# New opabiniid diversifies the weirdest wonders of the euarthropod lower stem group

**DOI:** 10.1101/2021.03.10.434726

**Authors:** Stephen Pates, Joanna M. Wolfe, Rudy Lerosey-Aubril, Allison C. Daley, Javier Ortega-Hernández

## Abstract

Once considered ‘weird wonders’ of the Cambrian, the emblematic Burgess Shale animals *Anomalocaris* and *Opabinia* are now recognized as lower stem-group euarthropods. *Anomalocaris* and its relatives (radiodonts) had a worldwide distribution and survived until at least the Devonian, whereas - despite intense study - *Opabinia* remains the only formally described opabiniid to date. Here we reinterpret a fossil from the Wheeler Formation of Utah as a new opabiniid, KUMIP 314087. By visualizing the sample of phylogenetic topologies in treespace, our results fortify support for the position of KUMIP 314087 beyond the nodal support traditionally applied. Our phylogenetic evidence expands opabiniids to multiple Cambrian Stages spanning approximately five million years. Our results underscore the power of treespace visualization for resolving imperfectly preserved fossils and expanding the known diversity and spatiotemporal ranges within the euarthropod lower stem group.

**Additional note:** This work contains a new biological name. New names in preprints are not considered available by the ICZN. To avoid ambiguity, the new biological name is not included in this preprint, and the specimen number (KUMIP 314087) is used as a placeholder.

Cover image.
Artistic reconstruction of the new opabiniid from the Wheeler Formation, Utah, USA (Cambrian: Drumian). Artwork by F. Anthony.

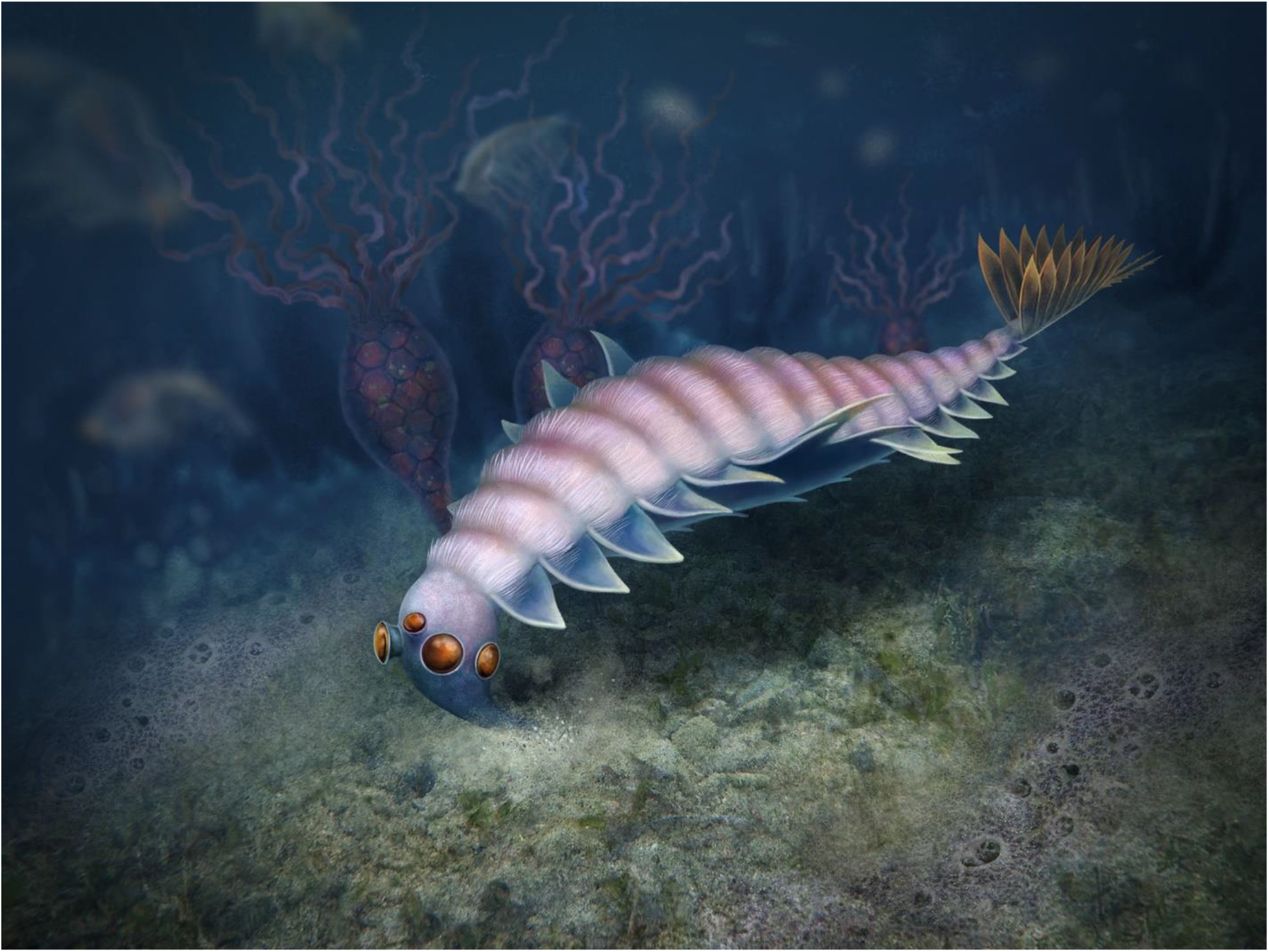

## Introduction

Euarthropods (e.g. chelicerates, myriapods, and pancrustaceans including insects) have conquered Earth’s biosphere, comprising over 80% of living animal species (Santos, de Almeida and Fernandes, 2021). Indeed, Euarthropoda has been the most diverse animal phylum for over half a billion years, documented by prolific trace and body fossil records that extend back to the early Cambrian (~537 and ~521 million years ago respectively) (Daley *et al.*, 2018). As the majority of these earliest euarthropods did not contain mineralised hard parts, we rely on remarkable fossil deposits such as the Burgess Shale, which preserve soft-bodied components of ancient biotas, to reveal critical data on the extraordinary diversity, disparity, and early evolution of Cambrian euarthropods (Gould, 1989).

Two of the most peculiar Burgess Shale animals, *Anomalocaris* and *Opabinia*, illustrate the complicated history of research of many Cambrian soft-bodied taxa - a result of their unfamiliar morphologies compared to the occupants of modern oceans (Collins, 1996; Briggs, 2015b). Both *Anomalocaris* and *Opabinia* possess compound eyes, lateral swimming flaps, filamentous setal structures, and a tail fan (Whittington, 1975; Whittington and Briggs, 1985; Budd and Daley, 2012; Daley and Edgecombe, 2014). Recent work has revealed that *Anomalocaris* and its relatives, the radiodonts, are united by the presence of paired sclerotized protocerebral frontal appendages and mouthparts composed of plates of multiple sizes, forming a diverse group containing over 20 taxa (Daley *et al.*, 2009; Cong *et al.*, 2014; Vinther *et al.*, 2014; Van Roy, Daley and Briggs, 2015; Ortega-Hernández, Janssen and Budd, 2017; Lerosey-Aubril and Pates, 2018; Liu *et al.*, 2018; Moysiuk and Caron, 2019). Radiodonts range in age from the early Cambrian to at least the Devonian, and have been recovered from numerous palaeocontinents (Kühl, Briggs and Rust, 2009; Daley and Budd, 2010; Cong *et al.*, 2014; Daley and Legg, 2015; Van Roy, Daley and Briggs, 2015; Fu *et al.*, 2019). Meanwhile, the most celebrated animal from the Burgess Shale (Budd, 1996; Briggs, 2015a), *Opabinia regalis*, with its head bearing five stalked eyes and a proboscis, remains the only opabiniid species confidently identified and is only known from a single quarry in the Burgess Shale. *Myoscolex ateles* from the Emu Bay Shale was proposed as a possible close relative (Briggs and Nedin, 1997), though this interpretation was hotly contested, and other authors have proposed a polychaete affinity (Glaessner, 1979; Dzik, 2004).

Radiodonts and *Opabinia* are now confidently placed within the lower stem of Euarthropoda (Budd, 1996; Daley *et al.*, 2009; Ortega-Hernández, 2016), following the assignment of nearly all Cambrian soft-bodied animals to stem and crown groups of modern phyla (e.g. Budd and Jensen, 2000). Fossils illustrating the sequence of character evolution along the euarthropod stem lineage provide the framework for understanding the evolutionary origins of the segmented, modular exoskeleton and the specialized appendages that underpin the ecological success of this phylum (Ortega-Hernández, 2016). Difficulties remain in interpreting the anatomical details, morphology, and phylogenetic placement of exceptional Cambrian fossils. In *Opabinia*, the presence of lobopodous limbs in addition to the swimming flaps cannot be confirmed, and the architecture of the flaps and associated setal blades remains elusive (Budd, 1996; Zhang and Briggs, 2007; Budd and Daley, 2012). Consequently, the phylogenetic position of *Opabinia* relative to radiodonts and deuteropods remains hotly debated. The identification of plesiomorphic and apomorphic characters has required new imaging and reinterpretations of existing specimens, the discovery of new fossil material and localities, and, crucially, the improvement of phylogenetic analysis methods to evaluate alternative relationships of enigmatic taxa.

Here we redescribe a fossil specimen from the Drumian Wheeler Formation of Utah, previously described as an anomalocaridid radiodont (Briggs *et al.*, 2008). KUMIP 314087 is a new genus and species that shares characters with both radiodonts and *Opabinia regalis*. We evaluate its phylogenetic position using both maximum parsimony (MP) and Bayesian inference (BI) and further interrogate the support for alternative relationships for KUMIP 314087 by visualizing the frequency and variation of these alternatives in treespace (Hillis, Heath and St. John, 2005; Wright and Lloyd, 2020). All analyses support an opabiniid affinity for KUMIP 314087. Our results evaluate the relative support for different hypotheses relating to the evolutionary acquisition of characters that define crown group euarthropods.

## Results

### Systematic Palaeontology

Superphylum PANARTHROPODA Nielsen, 1995

Family OPABINIIDAE Walcott, 1912

#### Diagnosis

Panarthropod with a short head region bearing a single unjointed appendage (‘proboscis’); slender trunk with dorsally transverse furrows delimiting segments; one pair of lateral flaps per body segment; setal blades cover at least part of anterior margin of lateral flaps; caudal fan composed of paired caudal blades; pair of short caudal rami with serrated adaxial margins.

#### Type genus

*Opabinia* Walcott, 1912.

#### Constituent taxa

KUMIP 314087 nov., *Opabinia regalis* Walcott, 1912.

#### Remarks

See **supplementary materials**.

> Genus *nov*.

#### Etymology

XXX

#### Type material, locality, and horizon

KUMIP 314087, part only, a complete specimen preserved compressed dorso-laterally. Collected from strata of the upper Wheeler Formation (Miaolingian: Drumian), at the Carpoid Quarry (GPS: 39.290417, −113.278519), southwest Antelope Mountain, House Range, Utah, USA (Briggs *et al.*, 2008).

#### Diagnosis

Opabiniid with slender trunk composed of at least 13, likely 15, segments; setal structures form blocks that cover the entire dorsal surface of the body and part of the anterior basal margin of the lateral flaps (setal blades only on flaps in *Opabinia*); tail fan composed of at least seven pairs of elongate caudal blades (three pairs in *Opabinia*).

> Genus and species *nov. gen. et sp*.
>
> Figs. 1b, c; 2b, d; 3
>
> *2008 Anomalocaris sp.: Briggs et al., p. 241, fig. 3*
>
> *2015 Anomalocaris sp.: Robison, Babcock and Gunther, p. 54-55, fig. 153 (top left)*
>
> *2012 Incertae sedis: Pates et al.,p. 29, table 1*.

**Figure 1.**
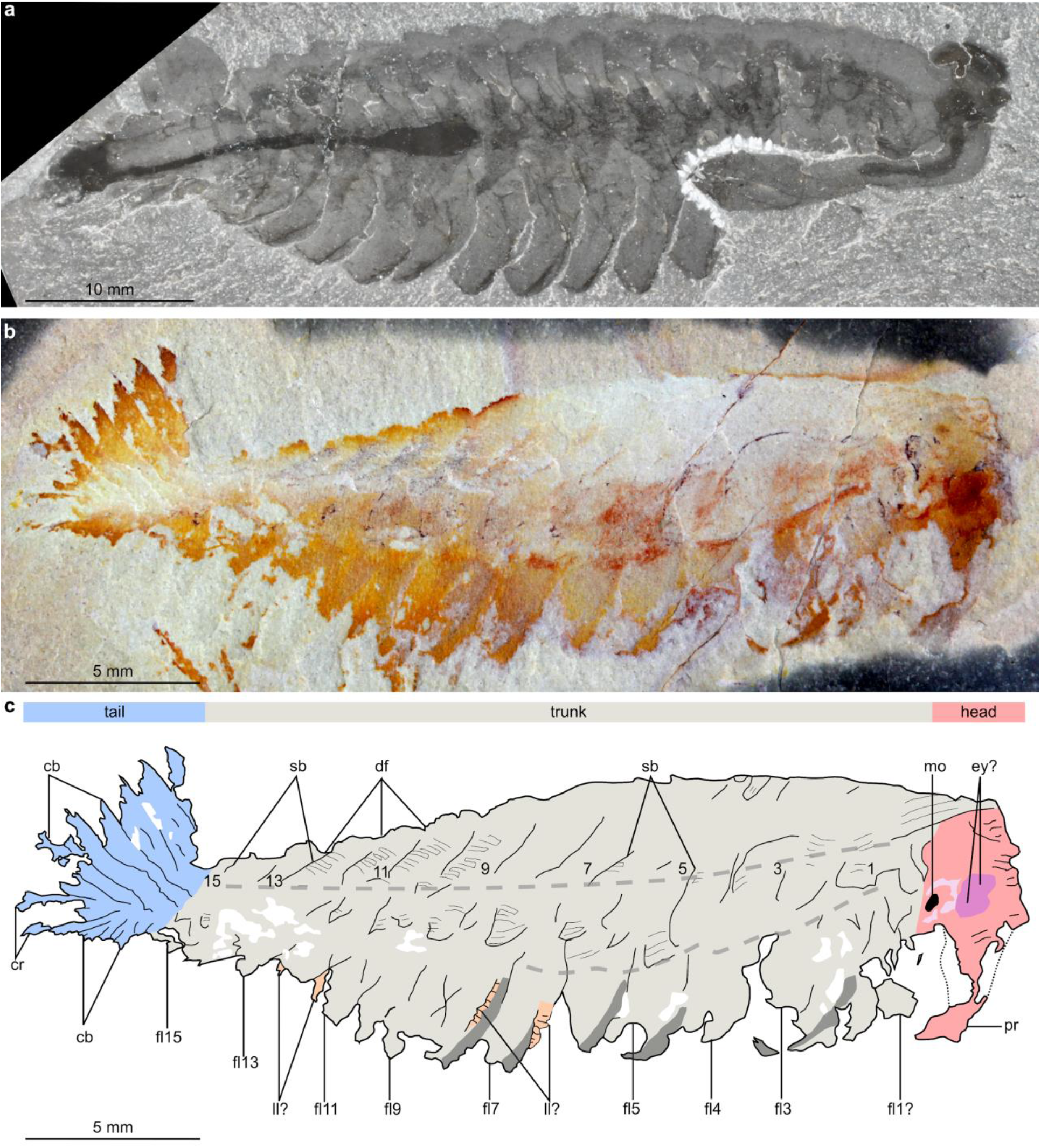
Comparison of *Opabinia regalis* Walcott, 1912 from the Burgess Shale (Cambrian: Wuliuan), British Columbia, Canada, and KUMIP 314087, gen. et sp. nov., from the Wheeler Formation (Cambrian: Drumian), House Range, Utah, USA. **(a)** USNM 155600, *Opabinia regalis* preserved in lateral view. **(b)** KUMIP 314087, preserved in dorsolateral view. **(c)** Interpretative drawing of panel B, dotted lines indicate inferred changes in slope on the body, numbers indicate body segments. Abbreviations: *cb*, caudal blade; *cr*, caudal ramus; *df*, dorsally transverse furrow delineating trunk segments; *ey?*, dark oval structure in head region, potential eye; *fl*, lateral flap; *ll?* potential lobopodous limb; *mo*, mouth; *pr*, proboscis; *sb*, setal blade block.

**Figure 2.**
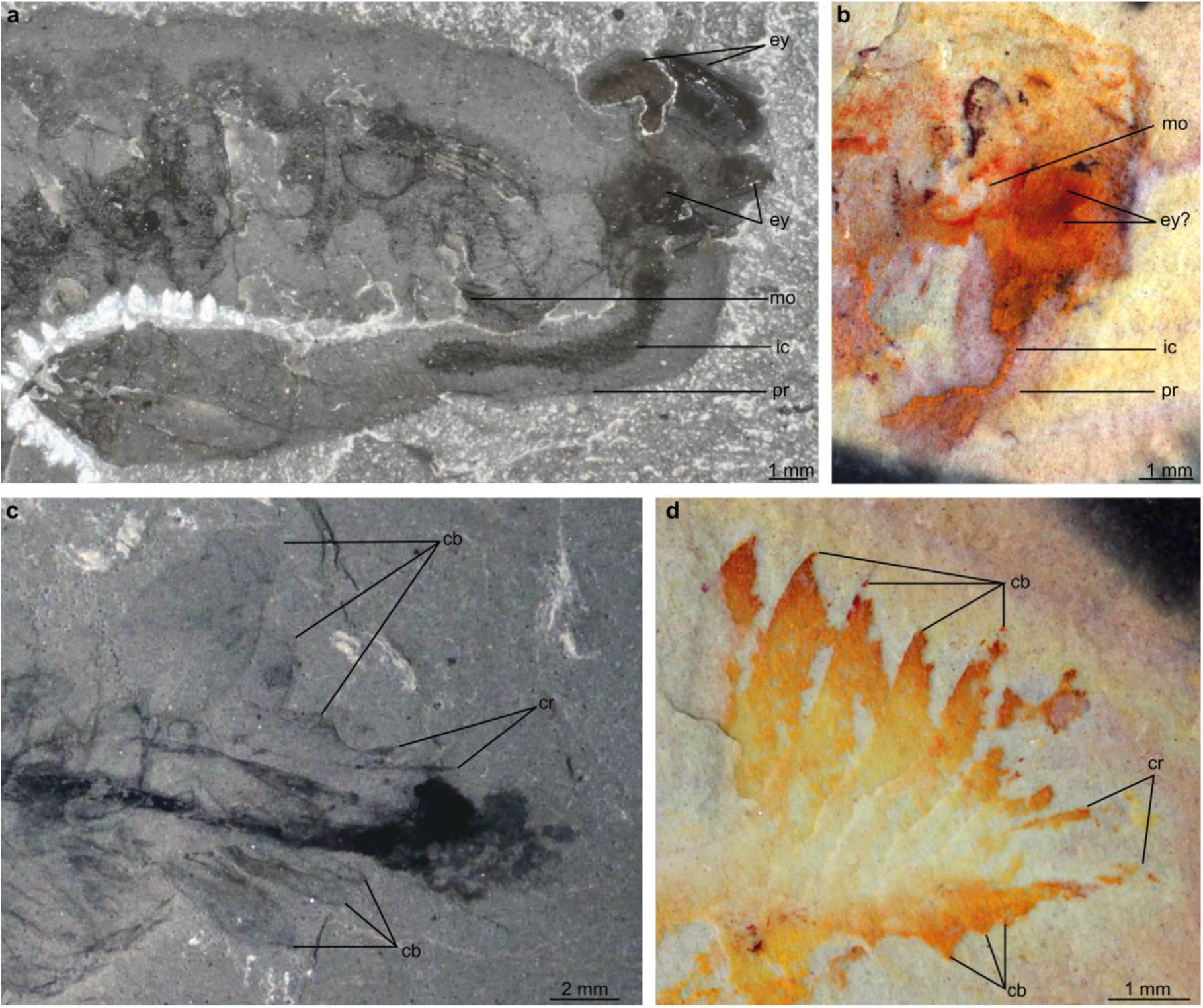
Details of the head and tail regions of *Opabinia regalis* Walcott, 1912 (a, c) and KUMIP 314087 gen. et sp. nov. (b, d). **(a)** Head region of USNM 155600, *Opabinia regalis*, showing eyes, posterior facing mouth, and proboscis with internal cavity. **(b)** Head region of KUMIP 314087, showing possible eyes, mouth, and putative proboscis with internal cavity. **(c)** Tail region of USNM 155600, *Opabinia regalis*, showing lobate tail blades, paired caudal rami with serrated adaxial margin, and posterior body termination extending beyond posteriormost caudal blades and caudal rami. **(d)** Tail region of KUMIP 314087 (photo mirrored), showing caudal blades and caudal rami with serrated adaxial margin. Abbreviations: *cb*, caudal blade; *cr*, caudal ramus; *ey*, eye; *ic*, internal cavity of proboscis; *mo*, mouth; *pr*, proboscis.

**Figure 3.**
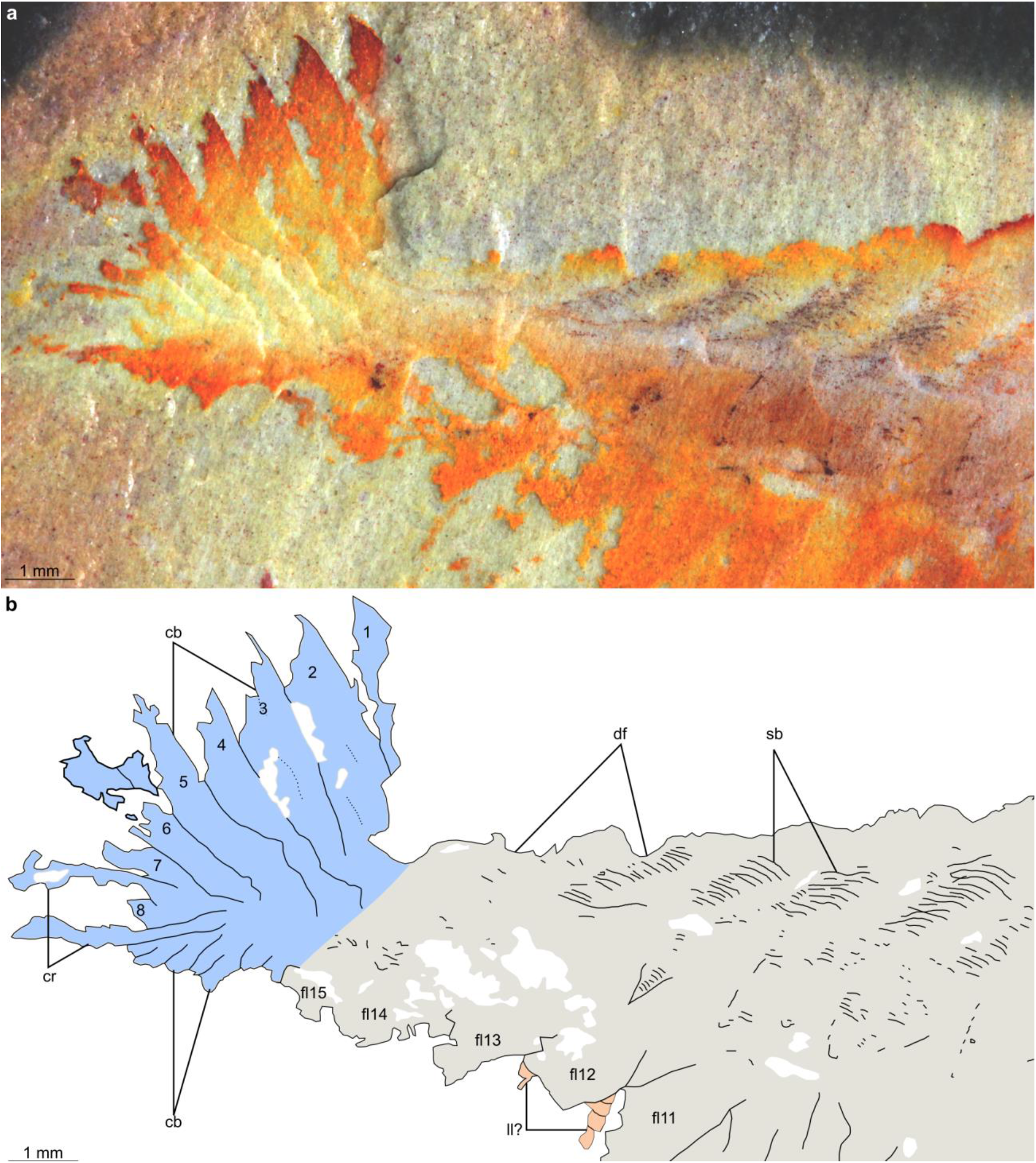
Posterior of KUMIP 314087 gen. et sp. nov. including details of setal blade blocks, elaborate tail fan, and paired caudal rami. **(a)** Photograph of specimen KUMIP 314097. **(b)** Interpretative drawing. Abbreviations: *cb*, caudal blade; *cr*, caudal ramus; *df*, dorsally transverse furrow delineating trunk segments; *fl*, lateral flap; *sb*, setal blade block.

**Table 1.** Number of trees bearing each bipartition of interest under different analytical regimes, with the total number of trees retrieved by each analysis in the last row. Categories are not mutually exclusive (e.g. *Opabinia* + Deuteropoda contains *Opabinia* only and Deuteropoda only trees), therefore the total of all rows exceeds the total number of trees in the analysis.

| **KUMIP 314087**<br>**forms a clade**<br>**with** | **Proboscis present** |  |  | **Proboscis uncertain** |  |  |
| --- | --- | --- | --- | --- | --- | --- |
|  | **Minimize**<br>**assumptions** | **Maximize**<br>**information** | **MP** | **Minimize**<br>**assumptions** | **Maximize**<br>**information** | **MP** |
| *Opabinia* | 1941 | 1149 | 12 | 1573 | 884 | 10 |
| Deuteropoda +<br>Radiodonta<br>(excludes<br>*Opabinia*) | 460 | 307 | 0 | 552 | 394 | 0 |
| *Opabinia* +<br>Deuteropoda | 359 | 105 | 12 | 401 | 104 | 12 |
| Deuteropoda | 186 | 65 | 0 | 225 | 73 | 2 |
| Radiodonta | 14 | 14 | 0 | 24 | 27 | 0 |
| *Pambdelurion* | 1 | 0 | 0 | 3 | 5 | 0 |
| *Kerygmachela* | 0 | 0 | 0 | 1 | 0 | 0 |
| Hurdiidae | 0 | 0 | 0 | 0 | 1 | 0 |
| None of the<br>above | 132 | 87 | 0 | 173 | 99 | 0 |
| **Total trees** | **2856** | **1656** | **12** | **2704** | **1512** | **12** |

#### Etymology

XXX

#### Diagnosis

As for genus, by monotypy.

#### Description

KUMIP 314087 represents a complete specimen preserved as a compression in dorsolateral view, with a length (sagittal) of 29 mm (**Fig. 1**). The overall organization consists of a short head, an elongate trunk with lateral body flaps, and a posterior tail fan.

The head region measures ~10% of the total body length, and preserves traces of eyes, the mouth and the proboscis. In the ventral posterior region of the head, two curved red structures surround a circular opening, interpreted as a mouth opening (**“mo” in Fig. 2b**). The mouth opening is immediately proximal to a dark red region of two overlapping oval shapes, tentatively interpreted as a pair of lateral eyes (**“ey?” in Fig. 2b**). Ventral to this, a cream-coloured elongated conical structure extends from the head ventrally (**“pb” in Fig. 2b**), with a sub-millimetric orange linear structure of variable width located along its midline (**“ic” in Fig. 2b**). This is tentatively identified as a proboscis with an internal cavity (**Fig. 2b**).

The slender trunk (~72% total body length) is widest towards the anterior and tapers towards the posterior. The dorsal margin bears a ‘corrugated’ appearance, with indents marking the point where dorsal intersegmental furrows intersect with the margin of the body (**“df” in Figs. 1, 3**). Blocks consisting of dozens of parallel darkly pigmented fine linear structures are arranged along the dorsal furrows and are interpreted as setal blades (**“sb” in Figs. 1, 3**). These blocks extend across the entire dorsal surface of the animal and continue laterally over the change in slope on the right side of the body. These setal structures display a triangular termination, which overlaps the anterior part of the base of the flaps (**Figs. 1, 3**).

At least 14, likely 15, of these lateral flaps are present on the right side of the body (**“fl1?-15” in Fig. 1**). Boundaries are not clear between what are interpreted as the two anteriormost flaps, and these may represent a single flap (**“fl1?” in Fig. 1**). Lateral flaps have a subtriangular outline and display a slight taper in size as the body thins posteriorly. The lateral flaps show reverse imbrication with the anterior margin of individual flaps overlapping the posterior margin of the flap immediately anterior to it. The surfaces of the flaps appear smooth and unornamented, with no evidence of strengthening rays or other internal features preserved, but the anterior margins of flaps 2-8 are preserved with a darker coloration compared to the inner region (**Figs. 1, 3**). Towards the posterior of the animal, a thin structure protruding from underneath a lateral flap could represent part of a ventral lobopodous limb, though the presence of additional material in the matrix of a similar width and orientation makes such an identification only very tentative (**“ll?” in Fig. 3**). Structures of a similar width can be seen related to flaps seven and eight (**“ll?” in Fig. 1**), though these may represent a poorly preserved anterior margin of these flaps.

The posterior of the body (~18% total body length) consists of a tail fan composed of paired elongate blades, and a pair of caudal rami. The tail has been twisted slightly and the right set of tail blades has been preserved flattened ventrally due to the dorsolateral aspect of preservation. The tail fan has seven, likely eight blades on the left side (**“cb” in Fig. 2**), while those on the right cannot be counted with certainty. Unlike the body flaps, these caudal blades are not associated with setal structures. They overlap one another proximally, a given blade largely concealing the blade immediately anterior to it. Each blade has the outline of an elongate parallelogram, longer on the anterior than posterior margin, and their acuminate distal regions splay out. The caudal rami are short (~3 mm length), converge towards a common point at the posterior of the animal, extend from the body at a different angle to the caudal blades, and exhibit serrated axial margins (**“cr” in Figs. 1, 2**).

### Remarks

KUMIP 314087 was originally described as an anomalocaridid radiodont based on the similarity in the shape of caudal blades to *Anomalocaris* taxa and the reverse imbrication of the flaps (Briggs *et al.*, 2008). KUMIP 314087 also shares with radiodonts the presence of setal blades that extend over the dorsal midline of the body. The recognition herein of a putative proboscis with internal cavity, dorsally transverse furrows that delimit segments in the trunk, and a short pair of caudal rami with serrated axial margins, support closer affinities of this animal with *Opabinia regalis*, rather than with *Anomalocaris*. The unique combination of characters, and novel features such as the elaborate tail fan, warrant the erection of a new genus and species.

Among members of the euarthropod lower stem-group, a proboscis has only been reported previously in *Opabinia* (Whittington, 1975). The proboscis of KUMIP 314087 protrudes from the head in a similar position relative to the tentatively interpreted eyes as in *Opabinia*. In addition, a feature comparable to the internal cavity within the proboscis of *Opabinia* can be observed in KUMIP 314087 (**Fig. 2**). However, unlike *Opabinia*, no annulations can be seen in this structure, as it is too poorly preserved. KUMIP 314087 also has dorsal furrows delineating the body segments. Such dorsal epidermal segmentation is seen in *Opabinia* but is unknown in all other lower stem group euarthropods (including *Kerygmachela, Pambdelurion* and all radiodonts) (Ortega-Hernández, 2016).

KUMIP 314087 also displays characters known in both radiodonts and *Opabinia*. The slender, broadly rectangular dorsal outline of the body in KUMIP 314087 is comparable to what is observed in both *Opabinia* and the radiodonts *Aegirocassis* and *Hurdia*. This outline contrasts with the diamond-like outline of many radiodonts, including *Amplectobelua symbrachiata, Anomalocaris canadensis*, and *Peytoia nathorsti* (Whittington and Briggs, 1985; Chen, Ramsköld and Zhou, 1994; Daley and Edgecombe, 2014). In addition, both *Opabinia* and radiodonts possess setal blades, in varying arrangements (**Supplementary Fig. 1**). In *Aegirocassis* and *Peytoia nathorsti*, these structures form a single block per body segment, which covers the entire dorsal surface (Van Roy, Daley and Briggs, 2015), while in *Opabinia* the setal structures cover the anterior margin of the flaps (Budd and Daley, 2012). KUMIP 314087 appears to display a combination of these two states, with setal blades covering the dorsal surface in a single block, which extends laterally to the basal region of the anterior margins of corresponding flaps (**Fig. 3**). Strengthened anterior margins of lateral flaps have also been reported in a juvenile specimen of the amplectobeluid radiodont *Lyrarapax* (Liu *et al.*, 2018). A tail fan associated with caudal rami is also known in both *Opabinia* and some radiodonts, though the number of blades known in KUMIP 314087 (at least seven, likely eight, on each side) by far exceeds what is known in either *Opabinia* (three) or any radiodont (ranging from zero to three). The acuminate tips of elongate caudal blades of KUMIP 314087 are most similar in morphology to those of *Anomalocaris*, and contrast to the more lobate caudal structures known in *Opabinia* and other radiodonts such as *Hurdia* (**Fig. 2**) (Whittington, 1975; Chen, Ramsköld and Zhou, 1994; Daley, Budd and Caron, 2013; Daley and Edgecombe, 2014). Paired caudal rami are also known in *Anomalocaris saron*, though these are much more elongate than in both KUMIP 314087 and *Opabinia* and lack the serrated adaxial margin common to the opabiniid taxa (**Fig. 2**) (Whittington, 1975; Chen, Ramsköld and Zhou, 1994).

#### Phylogenetic results

To test the affinities of KUMIP 314087 relative to *Opabinia* and radiodonts, we scored this specimen into a morphological matrix. Regardless of whether the matrix was analyzed with Bayesian inference (BI; **Fig. 4a, Supplementary Fig. 2a, 2b**) or maximum parsimony (MP; **Supplementary Fig. 2c**), a clade comprising KUMIP 314087 and *Opabinia* was resolved, warranting the assignment of the new taxon to family Opabiniidae. As the evidence for a proboscis in KUMIP 314087 is tentative (**Fig. 2b**), we conducted sensitivity analyses by building phylogenies where the proboscis (character 14) was coded as uncertain. With BI, opabiniids remained monophyletic (with lower nodal support; **Supplementary Figs. 3a, 3b**). With MP and an uncertain proboscis, the monophyly of opabiniids collapsed to a polytomy with deuteropods (**Supplementary Fig. 3c**).

**Figure 4.**
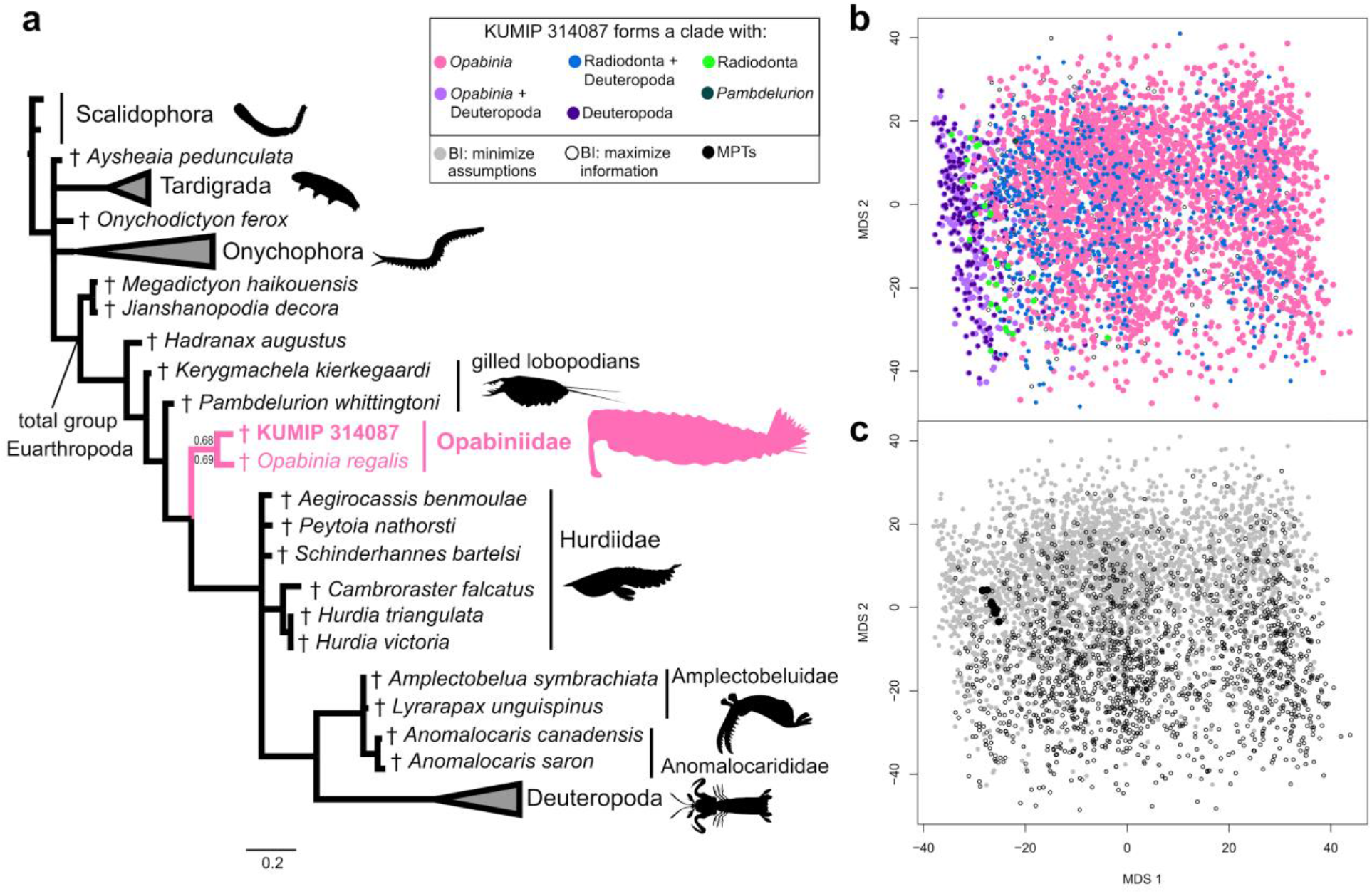
Phylogenetic relationships of opabiniids and lower stem group euarthropods. **(a)** Summarized topology based on the consensus tree retrieved with BI under minimize assumptions parameters. Numbers at the key node indicate posterior probabilities from this analysis, and from BI under maximize information parameters. Credits for silhouettes: *Priapulus caudatus*: Bruno C. Vellutini (CC BY 3.0); *Hurdia victoria* and *Anomalocaris canadensis*: Caleb M Brown (CC BY-SA 3.0) **(b)** Treespace plotted by bipartition resolving KUMIP 314087. Points are colored by relationships for this taxon. **(c)** Treespace plotted by analysis.

As the support values were poor for a morphological analysis (**Fig. 4a, Supplementary Fig. 2**: posterior probabilities of 0.68 and 0.69 with BI; jackknife value of 57 and GC value of 65 with MP), we visualized treespace (Hillis, Heath and St. John, 2005). Such plots identify whether uncertainty in support for opabiniid relationships in the posterior sample (*n* = 4512 trees for analyses where proboscis is coded as present; **Table 1**) and MPTs (*n* = 12 trees) is restricted to tree islands with otherwise similar topologies, or spread throughout a large region of occupied treespace. While treespace has been previously explored in meta-analyses of fossil datasets (Brazeau, Guillerme and Smith, 2019; Koch and Parry, 2020; Wright and Lloyd, 2020), this is, to our knowledge, the first attempt to use such a visualization to interrogate the distribution of bipartitions for the position of a focal fossil taxon. Several possible hypotheses are subsets: KUMIP 314087 could be part of a clade with either *Opabinia* or Deuteropoda (pink and dark purple colors, respectively, in **Fig. 4b**), and could be part of both those clades (light purple in **Fig. 4b**). Our overall treespace for KUMIP 314087 can nevertheless be grouped by islands of trees where the supermajority of trees are related to opabiniids (*n* = 3102 trees total for analyses where proboscis is coded as present) or a minority to deuteropods (*n* = 251 trees total). A sparse, slender zone (*n* = 28 trees total) of the alternative exclusive hypothesis that KUMIP 314087 is a radiodont (Briggs *et al.*, 2008) transitions between the opabiniid and deuteropod islands. Interspersed sparsely within the opabiniid island are topologies supporting KUMIP 314087 with both radiodonts and deuteropods, but excluding *Opabinia* (blue in **Fig. 4b**); most of these trees depict *Opabinia* as the direct outgroup rather than a wildcard taxon (occupying different positions that are topologically distant). Choice of BI model parameters did not substantially impact the treespace (**Fig. 4c**: grey and open circles overlap completely on axis 1 and much of axis 2), while the MPTs (**Fig. 4c**: black circles) formed a small but distinct cluster.

## Discussion

### The power of treespace for phylogenetic uncertainty of fossils

At first glance, our phylogenetic analyses provide only weak nodal support for the placement of KUMIP 314087 within Opabiniidae. Although similar nodal support with a similar data matrix has been used to reclassify enigmatic fossils (Howard *et al.*, 2020), we further interrogated our results - especially important as our terminal of interest is represented by a single specimen. Therefore, we investigated the degree of uncertainty in bipartitions, finding an increased number of topologies (**Table 1**) that support KUMIP 314087 related to at least one opabiniid, and not to an alternative taxon. Such calculations have been effective in summarizing the taxonomic uncertainty in fossil placement (Klopfstein and Spasojevic, 2019). Furthermore, our visualization of the sample of optimal trees (Hillis, Heath and St. John, 2005; St. John, 2017; Wright and Lloyd, 2020) illustrates the distribution of topological distances between conflicting and overlapping hypotheses. This technique allows the strength of support for competing hypotheses of relationships to be more comprehensively evaluated beyond an arbitrary cutoff value.

Phylogenetic analyses aiming to resolve the relationships of fossil taxa present challenges such as researcher-specific morphological interpretation and coding decisions, preponderance of missing data (common for exceptionally preserved Cambrian taxa, due to preservation of few specimens or taphonomic loss of labile morphology), and relatively simple models of character change that may not reflect true evolutionary history (Sansom, Gabbott and Purnell, 2010; Watanabe, 2016; Tarasov, 2019; Wright, 2019). Visualization of treespace investigates how these scenarios may affect a consensus topology. In the case of KUMIP 314087, the morphological description is based on a single specimen where we could only tentatively identify the proboscis. Therefore, we compared alternative codings to represent our uncertainty in interpretation, and the potential influence on the definition of opabiniids (**Supplementary Figs. 3, 4**). The sister group relationship of KUMIP 314087 with *Opabinia* (rather than radiodonts or deuteropods) is not driven by the proboscis character, and is maintained due to the other shared morphological characters (e.g. dorsal furrows, caudal rami).

### Implications for opabiniid evolution and ecology

Our phylogenetic results provide substantial support for an assignment of KUMIP 314087 to Opabiniidae, helping to clarify some debates about the morphology of *Opabinia*. Enigmatic triangular structures found along the body in *Opabinia*, have been variously interpreted as extensions of the gut (Whittington, 1975; Zhang and Briggs, 2007), or as lobopodous walking limbs (Budd, 1996; Budd and Daley, 2012). The potential lobopods extending from the ventral surface in KUMIP 314087 suggest that these walking limbs may be present in opabiniids generally. Additionally, two contrasting interpretations have been presented for the relationship between the lateral flaps and the blocks of setal blades in *Opabinia*: one where the setal blades are attached to the dorsal surface of the lateral flaps (Budd, 1996; Budd and Daley, 2012), and the other view suggesting the setal blades were attached as a fringe along the posterior margin of the lateral flap (Zhang and Briggs, 2007). The setal blades in KUMIP 314087 support the former interpretation, with the setal blades extending mainly along the dorsal surface of the body but also along the basal anterior margin of the flaps. Evidently the addition of even a single new specimen to the opabiniids provides crucial data informing on the group’s morphological aspects.

The family Opabiniidae is now considered to comprise two taxa, expanding its range geographically from two quarries separated by only a few meters to two deposits ~1000 km apart during two Cambrian Stages (Nanglu, Caron and Gaines, 2020). Although both *Opabinia* and *Anomalocaris* underwent major redescriptions around the same time (Whittington, 1975; Briggs, 1979; Whittington and Briggs, 1985), our revised opabiniids have not nearly caught up to the known diversity or distribution of radiodonts (or even the monophyletic groupings recovered in this study, Hurdiidae and Amplectobeluidae + Anomalocarididae). Radiodont frontal appendages, mouthparts, and carapaces are sclerotized and are often among the first fossils recovered from Cambrian deposits preserving non-biomineralizing organisms, and indeed many radiodont taxa are only known from their frontal appendages (e.g. Daley and Budd, 2010; Pates and Daley, 2019). However, preservation potential alone is insufficient to account for the greater diversity and distribution of radiodonts relative to opabiniids, as even radiodonts known only from complete specimens greatly outnumber opabiniids, both globally and within the Burgess Shale. Thus, the absence of opabiniids in other deposits from which complete radiodonts are known likely reflects a true absence or much lower diversity, which could have an ecological explanation.

Following a renaissance in radiodont research, it has been recognised that radiodonts display impressive variation in body size (milimetre to metre scale), body shape (rectangular to diamond shape in outline), inferred feeding ecology (raptorial predators, sediment sifters, filter feeders), and niche differentiation where species co-occur (e.g. Daley and Budd, 2010; Daley, Budd and Caron, 2013; Daley and Edgecombe, 2014; Vinther *et al.*, 2014; Van Roy, Daley and Briggs, 2015; Lerosey-Aubril and Pates, 2018; Liu *et al.*, 2018; Moysiuk and Caron, 2019; Lerosey-Aubril *et al.*, 2020; Pates *et al.*, 2021). In contrast, opabiniids show limited evidence for adaptations to different niches. Both taxa have rectangular body shapes and are centimetre scale (the single specimen KUMIP 314087 is ~50% the length of the largest *Opabinia*). The more elaborate tailfan of KUMIP 314087 may indicate a greater maneuverability of this taxon compared to *Opabinia regalis*, as the tailfan of *Anomalocaris canadensis* aided swift changes in direction (Sheppard, Rival and Caron, 2018).

### Implications for the euarthropod stem group

Our results have implications for larger scale questions, such as the relative phylogenetic positions of opabiniids and radiodonts along the euarthropod stem group, and detailed consideration of conflicting topologies. We replicate the dichotomy of recent publications, where matrices analyzed using MP find opabiniids as the sister group to deuteropods (Yang *et al.*, 2016, 2018; Howard *et al.*, 2020) and those analyzed using BI or maximum likelihood instead resolve radiodonts in that position (Fleming *et al.*, 2018; Moysiuk and Caron, 2019; Howard *et al.*, 2020; Zeng *et al.*, 2020). The branching order of these three clades has ramifications for the sequence of acquisition, and evolutionary reversals or convergences, of key crown group euarthropod characters (Ortega-Hernández, 2016), such as the posterior mouth and arthropodized appendages, as well as the dorsal expression of trunk segmentation (**Supplementary Fig. 5**). The scenario (favored by MP and an island of BI topologies) where opabiniids are sister group to deuteropods requires either the secondary loss of arthropodized appendages in opabiniids, or the convergent evolution of arthropodized appendages in radiodonts and deuteropods.

The consensus topology (**Fig. 4a, Supplementary Fig. 5a**), and the majority of topologies (yellow, pink, and maroon points in **Supplementary Fig. 5c**), support a single origin of arthropodization in euarthropods. A possible developmental framework would entail the single anterior protocerebral pair of arthropodized limbs in radiodonts becoming co-opted posteriorly to enable the arthropodization of all limbs (Jockusch, 2017; Chipman and Edgecombe, 2019). This scenario would require the convergent fusion of presumed protocerebral appendages in opabiniids to form a single proboscis, and of protocerebral limb buds in deuteropods to form the labrum (Chipman, 2015; Jockusch, 2017; Ortega-Hernández, Janssen and Budd, 2017; Park *et al.*, 2018). Evolutionary reversals or convergences are also required by these topologies (**Supplementary Figs. 5, 6**). The posterior-facing mouth shared by *Opabinia* and deuteropods is either convergent or lost in radiodonts (Ortega-Hernández, Janssen and Budd, 2017). Additionally, the distinct dorsally transverse furrows delineating segment boundaries (reported in both opabiniids), which may represent a precursor to arthrodized tergites in deuteropods (Yang *et al.*, 2015), could either be lost in radiodonts and regained in deuteropods, or represent a convergent expression of dorsal trunk segmentation.

The consensus topology is further complicated by the apparent paraphyly of radiodonts (**Fig. 4A, Supplementary Figs. 2a, 2b, 3b**). Traditional nodal support resolves a clade of amplectobeluids, anomalocaridids, and deuteropods with posterior probabilities of 0.52-0.61 (**Supplementary Figs. 2a, 2b, 3a, 3b**). The specific relationship of amplectobeluids and anomalocaridids with deuteropods might improve some aspects of limb evolution, as the loss of dorsal flaps (shared by opabiniids and hurdiids; **Supplementary Fig. 1**) prior to the proposed fusion of setal blades and ventral flaps into the deuteropod biramous limb removes the requirement to identify a dorsal flap homolog in deuteropods (Van Roy, Daley and Briggs, 2015). However, treespace visualization does not provide strong support for radiodont paraphyly, as overlapping islands resolve conflicting relationships among radiodonts and deuteropods (**Supplementary Figs. 5c, 7, supplementary discussion**). As many of the characters distinguishing internal relationships among radiodont families describe the protocerebral frontal appendages, and are coded as inapplicable to all other taxa, we propose revised models of character evolution (Tarasov, 2019; Wright, 2019) may be necessary to resolve these relationships; accordingly we place little weight on this particular result. It should be emphasized, however, that the position of KUMIP 314087 is not affected by this uncertainty, as its position as sister taxon to each radiodont clade was tested (with only non-zero results reported in **Table 1**).

### Conclusions

The “weird wonders”, as popularized by (Gould, 1989), inspired a generation of Cambrian paleontologists, with *Opabinia* at the heart of his narrative. The reorganization of previously enigmatic Cambrian taxa into stem groups instead revealed their importance for reconstructing the origins of modern phyla. Resolving the phylogenetic placement of these species is crucial for understanding the sequence of evolution of diagnostic crown group characters, as well as reconstructing the diversity and paleogeography of early ecosystems and groups. Here we apply treespace visualization to the reinterpretation of the relatively poorly preserved fossil KUMIP 314087. Dissection of the phylogenetic support demonstrates that while evidence for radiodont paraphyly is weak, KUMIP 314087 can be confidently reassigned to Opabiniidae. The weirdest wonder of the Cambrian no longer stands alone.

## Methods

### Fossil imaging and measurements

KUMIP 314087, accessioned at the Biodiversity Institute, University of Kansas, Lawrence, Kansas, USA (KUMIP), was photographed using a Canon EOS 500D digital SLR camera and Canon EF-S 60 mm Macro Lens, controlled for remote shooting using EOS Utility 2. Comparative figured material of *Opabinia regalis* is accessioned at the Smithsonian Institution U. S. National Museum of Natural History (USNM). Both polarized and unpolarized lighting were employed, with the fossil surface both wet and dry. Measurements were taken digitally using ImageJ2 (Rueden *et al.*, 2017).

### Morphological matrix

We added five fossil taxa (KUMIP 314087, *Amplectobelua symbrachiata* Hou, Bergström and Ahlberg 1995, *Anomalocaris saron* Hou, Bergström and Ahlberg 1995, *Cambroraster falcatus* Moysiuk and Caron 2019, and *Hurdia triangulata* Walcott 1912) and removed one fossil (‘Siberian Orsten tardigrade’) from a previously published morphological data matrix of panarthropods (Yang *et al.*, 2016), for a total of 43 fossil and 11 extant taxa. 86 characters were retained from the original matrix, 14 characters were added from two radiodont-focused datasets (Lerosey-Aubril and Pates, 2018; Moysiuk and Caron, 2019), and 25 characters were newly developed or substantially modified herein, for a total of 125 discrete morphological characters. Details of all characters including original and new character descriptions and scorings may be downloaded from MorphoBank (O’Leary and Kaufman, 2012) (www.morphobank.org, reviewer login ‘email address’: 3874, reviewer password: opabiniids).

### Phylogenetic analysis

The primary phylogenetic analyses were conducted using BI in MrBayes v.3.2.7 (Ronquist *et al.*, 2012), implementing the Markov (Mk) model (Lewis, 2001) of character change under two different parameter regimes. We followed the ‘maximize information’ and ‘minimize assumptions’ strategies of Bapst, Schreiber and Carlson (2018). The ‘maximize information’ strategy assumes equal rate distribution across characters and that state frequencies are in equilibrium, as in most previously published BI morphological studies. The ‘minimize assumptions’ strategy *(a)* applies gamma distributed among-character rate variation, and *(b)* varies the symmetric Dirichlet hyperprior with a uniform distribution of (0,10) to relax assumptions about character state frequency transitions (Wright, Lloyd and Hillis, 2016). As with complex molecular substitution models, the ‘minimize assumptions’ strategy may allow a better fit of the model to the data. Each analysis implemented four runs of four chains each (for 5.5 million and 9.5 million generations, respectively), with 25% burnin. Convergence was assessed based on standard deviations of split frequencies < 0.01, reaching effective sample size >200 for every parameter, and by comparing posterior distributions in Tracer v.1.7.1 (Rambaut *et al.*, 2018).

As the original matrix (Yang *et al.*, 2016) was devised for MP analysis, we explored MP topologies in TNT v.1.5 (Goloboff and Catalano, 2016) using implied weights (*k* = 3) and New Technology. We required the shortest tree to be retrieved 100 times, using tree bisection-reconnection to swap one branch at a time on the trees in memory (Wolfe and Hegna, 2014).

### Treespace analysis

Supplemental to traditional clade support metrics, we used classical multidimensional scaling (MDS) to plot treespace (Gower, 1966; Hillis, Heath and St. John, 2005; St. John, 2017; Wright and Lloyd, 2020), with the goal of identifying the distribution of trees resolving key clades formed with KUMIP 314087 (**Table 1**). Our R script inputs the unrooted post-burnin posterior samples (resultant from BI) and MPTs (resultant from MP) using *ape* v.5.3 (Paradis and Schliep, 2019), and employs *phangorn* v.2.5.5 (Schliep, 2011) to calculate pairwise unweighted Robinson-Foulds distances (RF, the proportion of bipartitions defined by a branch in one tree that is lacking in another tree) (Robinson and Foulds, 1981) for the total set of trees resulting from all analyses. The classical MDS function is performed on the RF distances, with a constant added to all elements in the distance matrix to correct for negative eigenvalues (Cailliez, 1983). The treespace therefore approximates the RF distances between trees (Hillis, Heath and St. John, 2005).

## Data availability

Supplementary data files are available at MorphoBank (www.morphobank.org, reviewer login (‘email address’): 3874, reviewer password: opabiniids) and at the Dryad Digital Repository, provisional link: https://datadrvad.org/stash/share/Vn0hPzM8ckUIiqlW9roY9dnMFi6qLOacYKiG1cs2GA8. Nomenclatural acts relating to the new taxon will be registered on ZooBank, LSIDXXX (publication), LSIDXXY (genus), LSIDXXZ (species).

## Acknowledgements

We are particularly grateful to P. Reese, who collected KUMIP 314087 and generously donated it to the Biodiversity Institute, University of Kansas (KUMIP). Access to and loan of this specimen was facilitated by B.S. Lieberman and J. Kimmig (KUMIP). J.O.H. thanks S. Whittaker (Smithsonian Institution) for facilitating access and training to imaging facilities. We thank M.J. Hopkins (American Museum of Natural History) and L.T. Rangel (Massachusetts Institute of Technology) for discussions about treespace, and F. Anthony for collaboration on the fossil reconstruction. S.P. acknowledges funding from an Alexander Agassiz Postdoctoral Fellowship (Museum of Comparative Zoology, Harvard University) and a Herchel Smith Postdoctoral Fellowship (University of Cambridge). This work was also supported by the National Science Foundation DEB #1856679 to J.M.W. and J.O.H.

## Author contributions

SP and JMW wrote the manuscript and created all the figures. All authors edited the text with critical insights. ACD, SP and JOH photographed fossil material. SP and JMW created the character matrix and coded taxa, with input from all other authors. SP led the fossil interpretation. JMW conceived and led the phylogenetic and treespace analyses.

## Extended Systematic Palaeontology

Family OPABINIIDAE Walcott, 1912

### Remarks

Walcott (1912) erected the Opabiniidae, a new family of anostracan crustaceans that included four genera: ‘ *Bidentia*’, *Leanchoilia*, *Opabinia*, and *Yohoia*. ‘ *Bidentia*’ is a junior synonym of *Leanchoilia* (Bruton and Whittington, 1983), which with *Yohoia* now belongs to the class Megacheira. *Opabinia* is therefore the only remaining representative of the family Opabiniidae. Hutchinson (1930) created a new anostracan suborder Palaeanostraca to group this family with the Rochdalidae and Yohoidae. None of the components of these families are regarded as related to anostracans anymore: *Opabinia* is considered a stem group euarthropod, *Rochdalia* is the nymph of an insect (Rolfe, 1967), *Yohoia* and *‘Bidentia’/Leanchoilia* are megacheirans (Hou and Bergström, 1997), and *Branchipusites* was reinterpreted as an arthropleurid (Guthörl, 1934). Accordingly, we regard the suborder Palaeanostraca as invalid. More recently, Collins (1996) placed Opabiniidae into a new class, Dinocarida, along with members of the order Radiodonta. This clade has been shown to be paraphyletic in several phylogenetic analyses (e.g. Daley *et al.*, 2009; Van Roy, Daley and Briggs, 2015; Lerosey-Aubril and Pates, 2018).

The genus *Opabinia* originally included two species, *Opabinia regalis* (type species) and *Opabinia? media*. Hutchinson (1930) considered the latter species to be composed of juvenile of *O. regalis*, and therefore to represent a junior synonym of this species. By contrast, Simonetta (1970) and Whittington (1975) regarded *Opabinia? media* as an invalid taxon, the three specimens putatively used by Walcott (1912) to define this species being recognized as belonging to a different genus or considered too poorly preserved to be assigned to any known taxon. A poorly preserved fossil recovered from the Furongian of Siberia was later used to describe a new species, *Opabinia norilica* (Miroshnikov and Krawzov, 1960), but this was rejected by Whittington (1975). In summary, *Opabinia* is composed solely of the type species, *Opabinia regalis*.

In this study we find phylogenetic support for including a new taxon KUMIP 314087 in the family Opabiniidae, under both maximum parsimony and Bayesian inference methods. Morphological support for uniting these taxa comes from a shared proboscis (not well preserved in KUMIP 314087), dorsal intersegmental furrows in the trunk, and small lateral flaps. KUMIP 314087 and *Opabinia regalis* also both possess a tail fan with paired caudal rami, and setal structures which cover the anterior of the flap, though those of KUMIP 314087 also cover the entire dorsal surface. The presence of 15 flap-bearing trunk segments might also be common to the two taxa (though the exact number is only tentative in KUMIP 314087, and this character is not included in the phylogenetic analysis). *Opabinia regalis* has stalked eyes, but while eyes are identified in the head region of KUMIP 314087, there is no evidence that these structures are stalked.

Notably, while a presence of a proboscis, perhaps the most famous and unusual character observed in *Opabinia regalis*, is included in the diagnosis for the family, it may be possible to identify future opabiniids lacking this feature. The proboscis is known in *Opabinia* and tentatively in KUMIP 314087. Our phylogenetic analyses where a proboscis is coded as uncertain in KUMIP 314087 also return this taxon as sister group to *Opabinia* (**Supplementary Fig. 3A, B, 4**) and therefore as a member of Opabiniidae.

### Additional implications of our phylogenetic results

#### Radiodonta paraphyly?

Using MP, Radiodonta is resolved as a monophyletic group, alongside a very small percentage of BI posterior trees (teal points in **Supplementary Fig. 7**). The majority of BI posterior trees, and therefore the majority of the treespace, support radiodont paraphyly (**Supplementary Fig. 7**).

When the trees are separated by what is sister to deuteropods, around half the space is occupied by trees supporting a monophyletic Deuteropoda + Amplectobeluidae + Anomalocarididae, however the clade Amplectobeluidae + Anomalocarididae can be either monophyletic or paraphyletic in this plot (**Supplementary Fig. 5**). When this area of the treespace is compared to what is occupied by monophyletic Amplectobeluidae + Anomalocarididae (**Supplementary Fig. 7**), there is only limited overlap in these two areas, lowering the support for the tree topology depicted in the consensus.

There is evidence that internal radiodont relationships are not confidently resolved based on the data (character matrix) and/or model of morphological character change. For example, trees where the family Hurdiidae is recovered as monophyletic are spread across the whole treespace, but at a low density. Often a tree with monophyletic Hurdiidae is very close in the treespace (i.e. with a low RF value or similar overall topology) to multiple trees where Hurdiidae is not resolved as a monophyletic group (**Supplementary Fig. 7**). Future work (ongoing) aimed at better resolving radiodont internal relationships, as well as improving model and matrix design to deal with this kind of problem common to palaeontological datasets will allow more certainty to be placed on the monophyly or paraphyly of radiodonts. Little weight should be placed on this particular result from this study, as the support for radiodont monophyly is poor and not enhanced by additional bipartitions (as in opabiniids), and as other matrices consistently resolve radiodonts as a monophyletic group when analysed with both MP and BI (e.g. Lerosey-Aubril and Pates, 2018; Moysiuk and Caron, 2019; Zeng *et al.*, 2020).

## Supplementary Figures

**Supplementary Figure 1.**
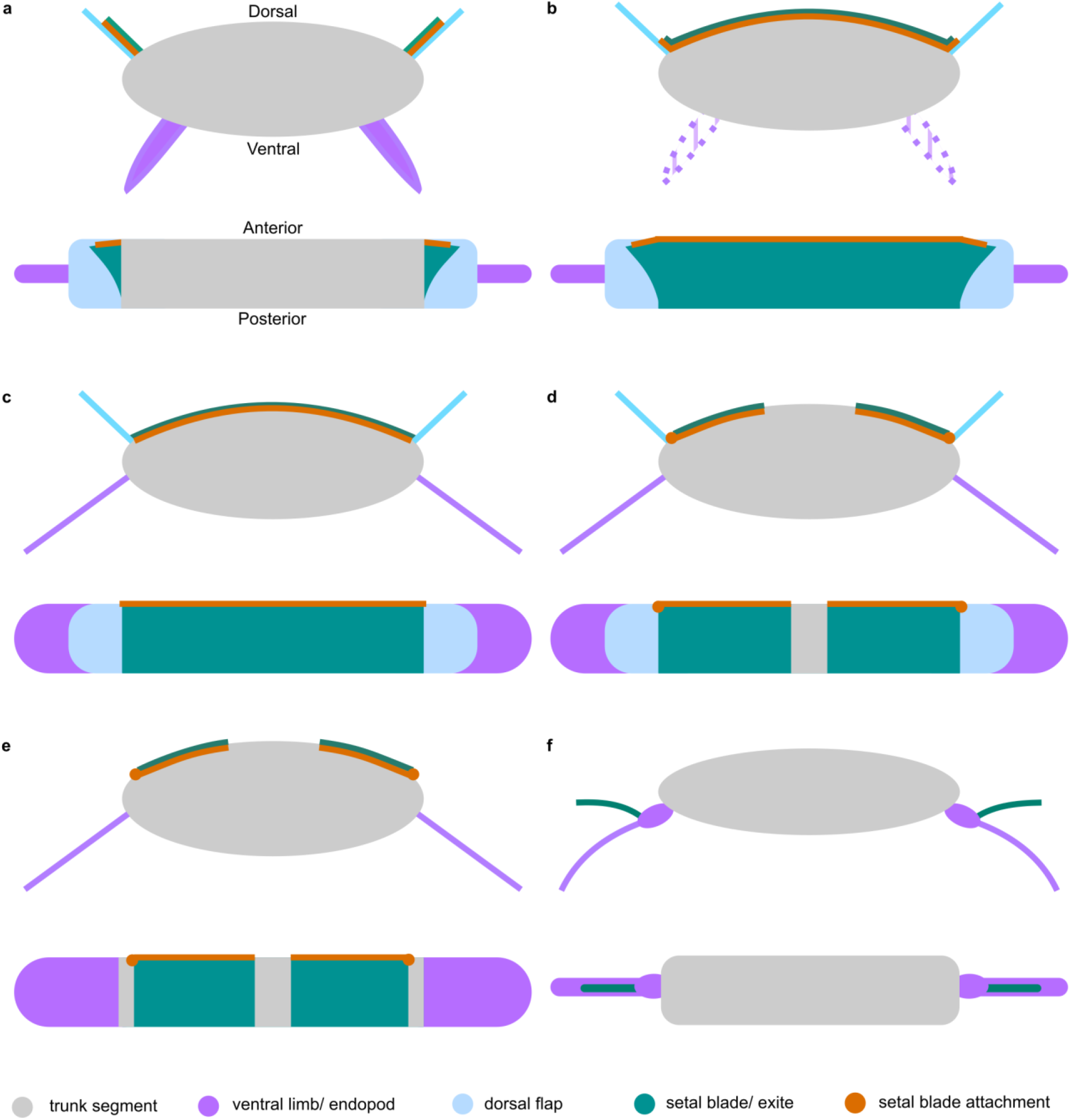
Comparison of setal blade attachment in opabiniids and radiodonts, showing current hypothesis of homology to the biramous limb of deuteropods (Van Roy, Daley and Briggs, 2015). **(a)** *Opabinia regalis*. **(b)** KUMIP 314087. **(c)** *Aegirocassis benmoulai, Peytoia nathorsti*. **(d)** *Hurdia* spp. **(e)** *Anomalocaris* spp. **(f)** Generalized deuteropod.

**Supplementary Figure 2.**
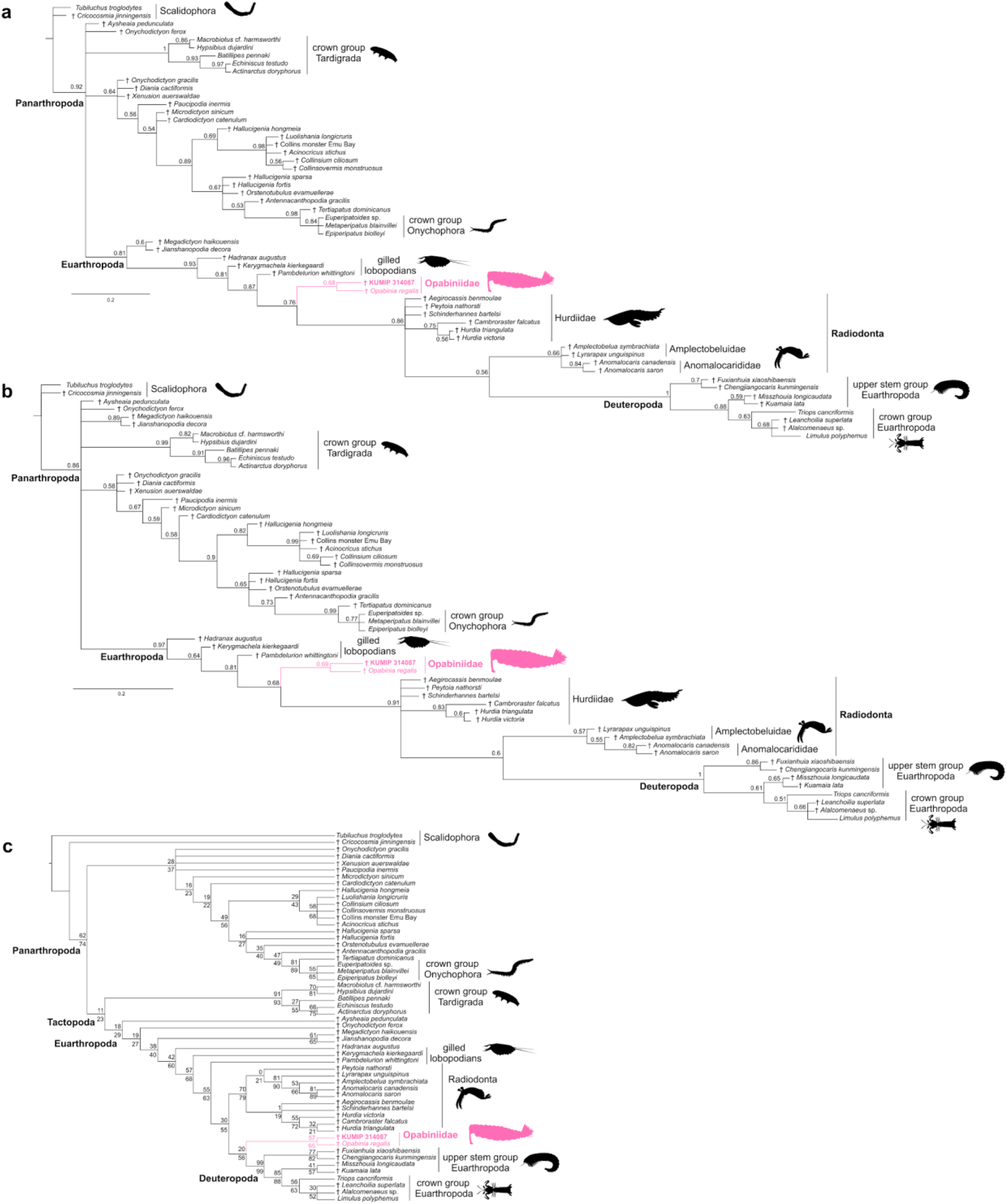
Full phylogenetic results for opabiniids. **(a)** Majority rule consensus tree retrieved with BI, under minimize assumptions parameters. Numbers above nodes indicate posterior probabilities. **(b)** Majority rule consensus tree retrieved with BI, under maximize information parameters. Numbers above nodes indicate posterior probabilities. **(c)** Strict consensus tree retrieved with MP (12 MPTs, 230 steps, consistency index: 0.687, retention index: 0.889). Numbers above nodes indicate jackknife values; numbers below nodes indicate group present/contradicted (GC) values from symmetric resampling.

**Supplementary Figure 3.**
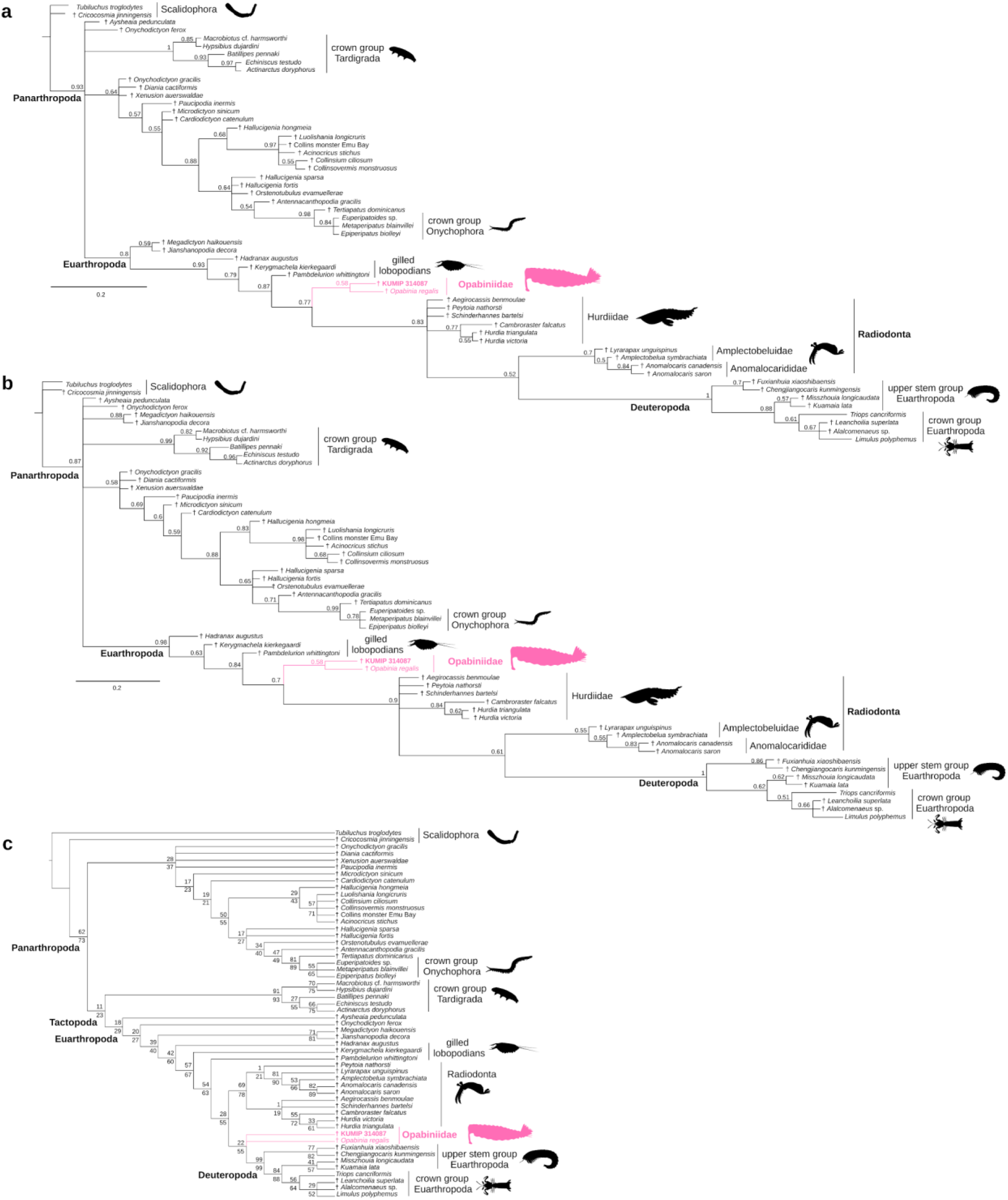
Full phylogenetic results for opabiniids, where proboscis is coded as uncertain in KUMIP 314087. **(a)** Majority rule consensus tree retrieved with BI, under minimize assumptions parameters. Numbers above nodes indicate posterior probabilities. **(b)** Majority rule consensus tree retrieved with BI, under maximize information parameters. Numbers above nodes indicate posterior probabilities. **(c)** Strict consensus tree retrieved with MP (12 MPTs, 230 steps, consistency index: 0.687, retention index: 0.888). Numbers above nodes indicate jackknife values; numbers below nodes indicate GC values from symmetric resampling.

**Supplementary Figure 4.**
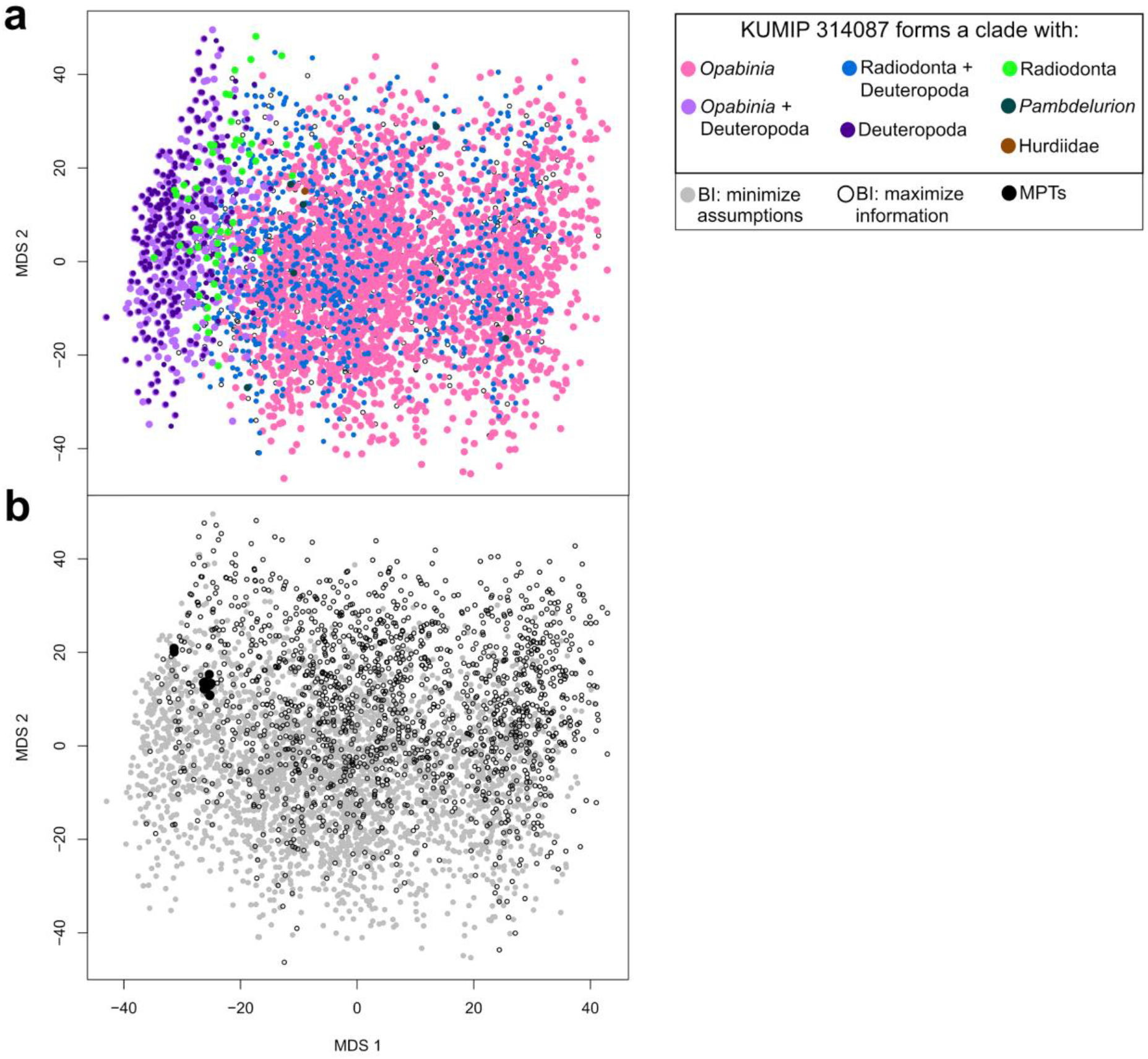
Treespace analysis for the modified matrix where KUMIP 314087 is recoded to reflect uncertainty in the presence of the proboscis. **(a)** Treespace plotted by bipartition resolving KUMIP 314087. Points are colored by relationships for this taxon. **(b)** Treespace plotted by analysis.

**Supplementary Figure 5.**
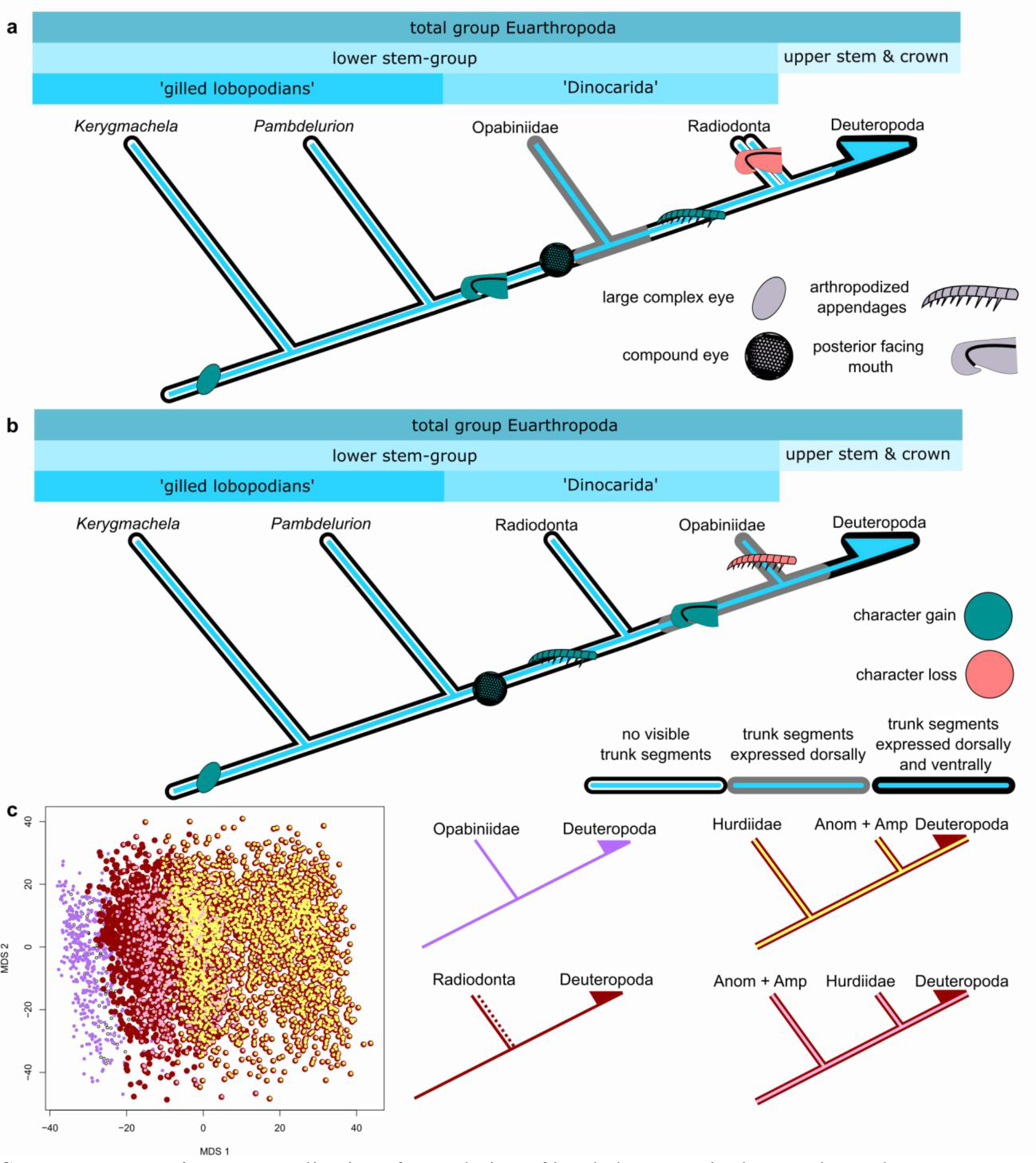
Implications for evolution of head characters in the euarthropod stem group, based on our phylogenetic results, considering only evolutionary reversals. **(a)** Topology resulting from all BI consensus trees. **(b)** Topology resulting from MP consensus trees. **(c)** Treespace plotted by bipartition resolving the sister group of Deuteropoda. Points are colored by relationships for this taxon.

**Supplementary Figure 6.**
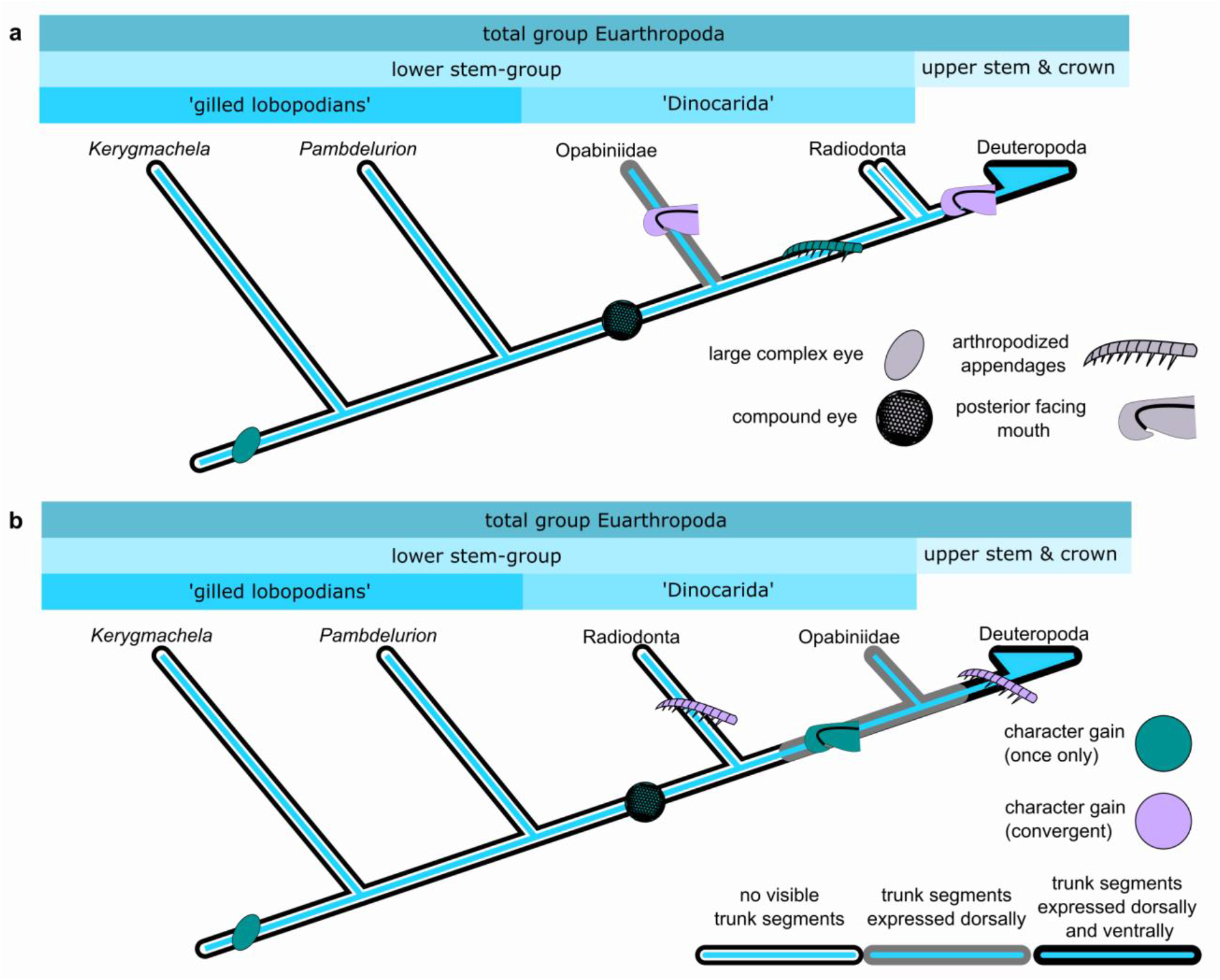
Implications for evolution of head characters in the euarthropod stem group, based on our phylogenetic results, considering only evolutionary convergences. **(a)** Topology resulting from all BI consensus trees. **(b)** Topology resulting from MP consensus trees.

**Supplementary Figure 7.**
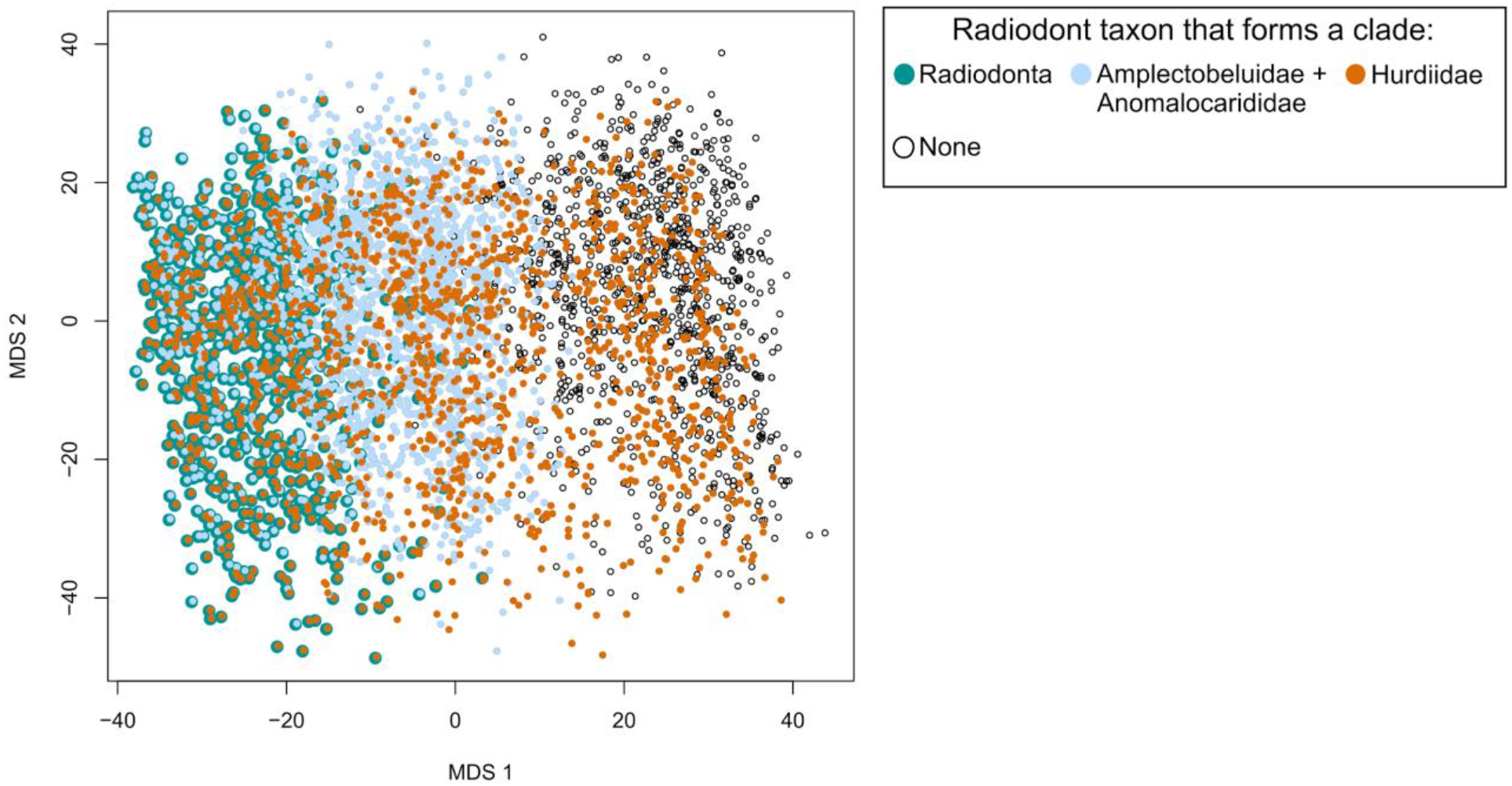
Treespace plotted by bipartition resolving the monophyly of radiodont taxa. Points are colored by relationships for these selected taxa.

